# Optimization of conditions for production of soluble *E. coli* polyA-polymerase

**DOI:** 10.1101/2024.12.01.626206

**Authors:** Igor P. Oscorbin, Maria S. Kunova, Maxim L. Filipenko

**Affiliations:** Institute of Chemical Biology and Fundamental Medicine, Siberian Branch of the Russian Academy of Sciences (ICBFM SB RAS), 8, Lavrentiev Avenue, Novosibirsk, 630090, Russia

**Keywords:** PcnB, polyA-polymerase, PAP-1, polyadenylation, E. coli

## Abstract

Poly(A)-polymerase (PAP-1) from *Escherichia coli* is the primary enzyme responsible for synthesizing poly(A) tails on RNA molecules, signaling RNA degradation in bacterial cells. In vitro, PAP-1 is used to prepare libraries for RNAseq and to produce mRNA vaccines. However, *E. coli* PAP-1’s toxicity and instability in low-salt buffers complicate its expression and purification. Here, we optimized the conditions for the production of recombinant PAP-1. For that, *E. coli* PAP-1 was expressed in seven *E. coli* strains with different origin and genetic backgrounds, followed by assessment of the overall protein yield, solubility, and enzymatic activity. Among the tested strains, BL21 (DE3) pLysS achieved the best balance of cell density, total PAP-1 yield, solubility, and specific activity. Rosetta 2 (DE3) and Rosetta Blue (DE3) hosting the pRARE plasmid exhibited the lowest solubility, likely due to excessive translation efficiency. Higher induction temperatures (>18°C) exacerbated PAP-1 insolubility. Interestingly, PAP-1 accumulation correlated with an increase in the plasmid copy number encoding the enzyme, indicating its potential utility as a surrogate marker for PAP-1 activity. These findings provide insights into optimizing *E. coli* PAP-1 production for biotechnological applications.

## Introduction

Polyadenylation is a well-known nuclear process in eukaryotic cells, but polyadenylated RNA molecules have also been found in bacteria. While the relationship between polyadenylation and RNA degradation has been investigated, the exact details of this cellular process remain unclear [1]. At least three bacterial enzymes — polynucleotidylphosphorylase (PNPase) [2], RNase PH [3], and poly(A) polymerase (PAP) — are known to synthesize polyadenylic tails in a template-independent manner. Among these, poly(A) polymerase is primarily responsible for most polyadenylation under normal conditions, and its malfunction leads to a significant reduction in polyadenylated RNA molecules [4–7]. PAP belongs to the nucleotidyltransferase superfamily, along with tRNA nucleotidyltransferases, which add CCA tails to tRNA molecules during tRNA maturation [8].

Despite its discovery in the early 1960s, only a few bacterial PAPs have been purified and biochemically characterized. These include two enzymes from *E. coli* [9,10], one from *Geobacter sulfurreducens* [11], and another from *Pseudomonas putida* [12]. However, the *Pseudomonas putida* PAP has not been cloned and might represent a different enzyme capable of synthesizing poly(A) tails. Among these, *E. coli* PAP-1 is the most extensively studied, with optimal conditions for temperature, cofactors, salt concentration, and pH determined [13,14]. The pcnB gene encoding *E. coli* PAP-1 was identified as a determinant of plasmid copy number in ColE1 origins [4,15]. Later, Sarkar et al. defined the pcnB product as poly(A) polymerase [5], which had already been purified and characterized at that time. However, the biochemical properties of other bacterial PAPs have not been as thoroughly evaluated, limiting understanding of their cellular roles and potential practical applications. Recently, pcnB genes have been identified in multiple bacterial species, but their respective PAPs have not been cloned, and their properties remain unknown. This lack of empirical data also hampers the in silico prediction of pcnB genes and their differentiation from similar *cca* genes [16].

PAPs have gained attention as tools for polyadenylating in vitro synthesized RNA molecules, particularly libraries for RNAseq and mRNA for vaccine use. During the SARS-CoV-2 pandemic, mRNA vaccines demonstrated their potential to control viral spread, proving the technology’s promise for other applications such as anticancer vaccines [17,18]. As the development of mRNA vaccines accelerates, the demand for simplified and cost-effective reagents for their production increases. However, *E. coli* PAP-1 is known for its instability, including a tendency to lose activity due to slow aggregation in low-salt buffers [5,13]. Additionally, it contains eight cysteine residues, making it prone to remaining in insoluble fractions after cell lysis. Furthermore, *E. coli* tightly regulates PAP-1 expression because its overproduction is toxic to host cells, significantly reducing cell viability post-induction [6]. Together, these issues pose challenges for developing protocols to obtain soluble *E. coli* PAP-1 in high quantities.

One common approach to optimize recombinant protein production is selecting the most suitable host strain based on the protein’s solubility and yield [19,20]. Various *E. coli* strains have been engineered to improve recombinant protein production. Rosetta strains carry the plasmid pRARE, which encodes tRNAs rare in *E. coli*, enhancing the expression of heterologous proteins [21–23]. BL21 Gen-X is designed to increase expression yields, while SoluBL21 enhances the solubility of mammalian proteins [24–26]. Lemo21 (DE3) includes a plasmid encoding the T7 RNA polymerase inhibitor LysY, enabling expression tuning via L-rhamnose concentration [27,28]. Shuffle T7 cells express DsbC disulfide bond isomerase to correct non-native disulfide bonds, along with an additional LacI protein to prevent promoter leakage [29,30].

In this study, we aimed to identify the most effective *E. coli* strain for producing recombinant *E. coli* PAP-1. Seven *E. coli* strains were tested to evaluate the enzyme’s yield and solubility. Additionally, we measured the ratio of plasmid carrying the pcnB gene to the host cell’s genomic DNA and compared these ratios with the yield, solubility, and activity of the recombinant enzyme.

## Materials and Methods

### Cloning of *E. coli* polyA polymerase

The coding sequence of *E. coli* PAP-1 (GenBank: M20574.1) was amplified using PcnB- Eco-F1/PcnB-Eco-R1 primers (Table 1) with NheI and XhoI restriction sites, allowing the in-frame ligation into the pET36b vector (Novagen, Madison, WI, USA). PCR was carried out using genomic DNA of *E. coli* strain XL1-Blue as a template. The resultant 1.6-kbp DNA fragment and pET36b vector were digested with NdeI and XhoI (SibEnzyme, Russia), ligated, and transformed into *E. coli* XL1-Blue cells according to the standard protocols [31]. The fidelity of the resulting recombinant plasmid named pET-PAP-Eco was confirmed by sequence analysis using primers pET-F and pET-R (Table 1).

**Table 1.**
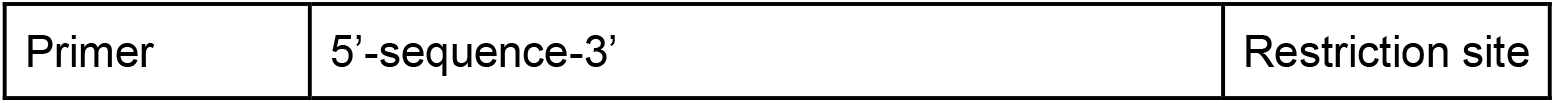
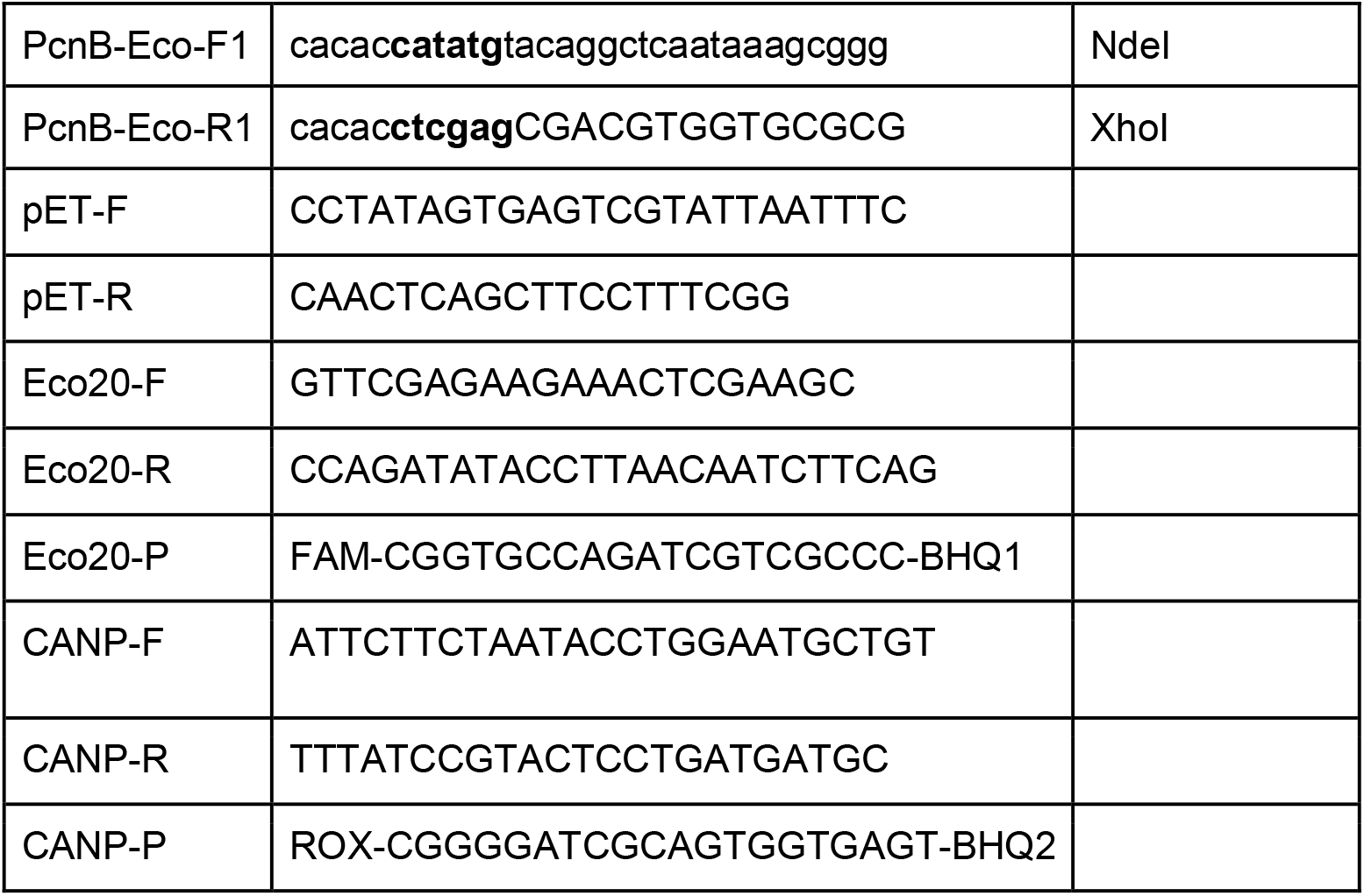
Oligonucleotide primers and probes.

### Expression of *E. coli* polyA polymerase

A starter culture of *E. coli* strains (Table 2) harboring the plasmid pET-PAP-Eco was grown to OD600 = 0.6 in LB medium with 25 μg/mL kanamycin at 37 °C. For each strain, two cultures in 5 mL of medium each were prepared. After reaching the designated optical density, probes for PAGE were taken as controls before induction, IPTG was added to a final concentration 1 mM to induce the expression of PAP-1, the cultures were divided to 1.5 mL subcultures that were subjected to ON incubation at 18, 25, or 37°C. After induction, the OD600 was measured, control probes after induction were sampled and cells from 1 mL were harvested by centrifugation at 4,000 × g and stored at −70 °C for possible solubility test.

**Table 2.**
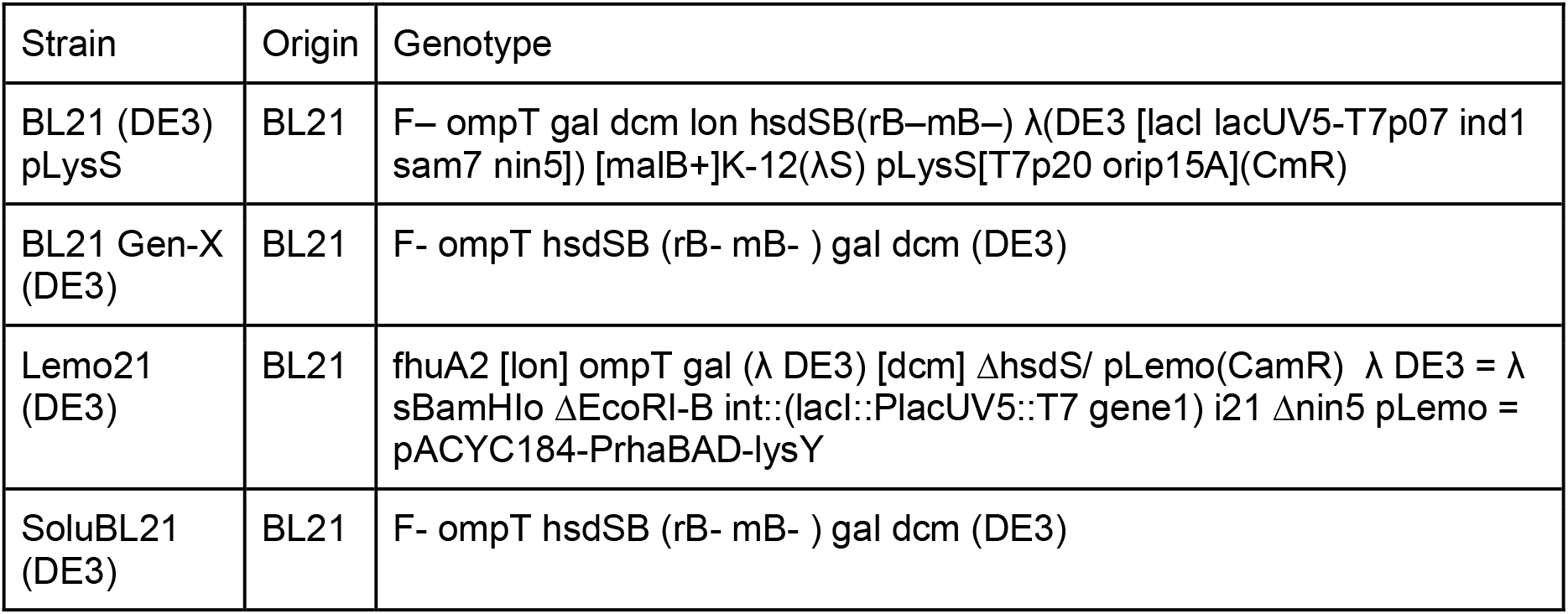
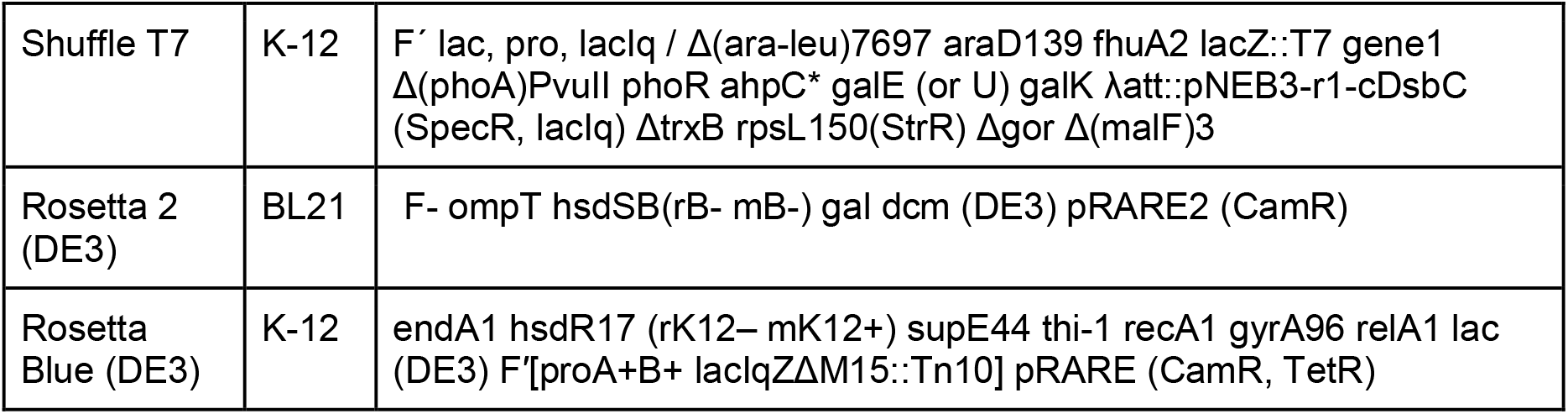
*E. coli* strains used for expression of PAP-1 *E. coli*.

### Solubility test

To evaluate solubility of PAP-1 *E. coli* in different E.coli strains, cell pellets from 1 mL night cultures were lysed in 200 μl of a lysis buffer containing 50 mM Tris-HCl, pH 8.0, 200 mM KCl, 1 mM ????, 5 mM beta-mercaptoethanol, 5% glycerol, 1 mM PMSF, 0.5% 3-(N,N- Dimethylmyristyl-ammonio)propanesulfonate, 0.05% C7BzO. The cell pellets were resuspended in the lysis buffer following addition of lysozyme to a 0.5 mg/ml and incubation for 1 hour at 37 °C. The incubated probes were sonicated and centrifuged at 20,000 × g for 15 min. Soluble fractions were transferred into new tubes, and insoluble pellets were resuspended in 200 μl of the lysis buffer. Both soluble and insoluble fractions were analyzed using SDS-PAGE in 12.5 % acrylamide gel, the resulting electropherograms were quantified by the ImageLab software (Bio-Rad, Hercules, CA, USA).

### Polyadenylation assay

Specific activity of poply(A) polymerase was analyzed using elongation of fluorescently labeled (r)A20 oligonucleotide. Reaction mixes contained in a volume of 10 μL 1× reaction buffer for PAP-1 (50 mM Tris-HCl, 250 mM NaCl, 10 mM MgCl2, pH 8.0), 1 mM ATP, 10 pmol of (r)A20- oligonucleotide, and a specified below amount of bacterial lysates after induction. The reactions were started by the addition of the enzyme and immediately transferred to a preheated thermocycler followed by an incubation for 30 minutes at 37 °C. After incubation, the reactions were quenched by addition of 10 μL of formamide and denatured by heating for 5 minutes at 95 °C. Reaction products were analyzed using denaturing PAGE in a 18% acrylamide gel with 7 M urea and quantified by the ImageLab software (Bio-Rad, Hercules, CA, USA).

### Quantitative PCR

Quantitative PCR reactions were performed in 20 µL volume containing 65 mM Tris-HCl, pH 8.9, 24 mM (NH4)2SO4, 0.05% Tween-20, 3 mM MgSO4, 0.2 mM dNTPs, 600 nM primers Eco20-F, Eco20-R, CANP-F, CANP-R, 100 nM TaqMan probes Eco20-P, CANP-P (Table 1) and 1 U of Taq-polymerase (Biosan, Novosibirsk, Russia). Amplification was carried out in CFX96 Real-Time PCR Detection System (Bio-Rad, Hercules, CA, USA) according to the following program: 95 °C for 3 min followed by 45 cycles of 95°C for 10 s, and 60°C for 40 s with a collection of fluorescent signals at FAM and ROX channels. The acquired data were analyzed with CFX Manager software (Bio-Rad, Hercules, CA, USA).

### Data analysis

Relative amount of the polyadenylated substrate was calculated as follows — Ratio = I(elongated products)/I(substrate)+I(elongated products)) — and used to estimate by a linear regression model a lysate amount necessary to elongate 50% of the substrate initially added to the reaction. These computed lysate volumes served to assess PAP-1 activity. To account for a final cell density, the computed lysate volumes were divided on the respective OD600 values.

Correlations between variables (induction temperature, OD600, PAP-1 level, etc) were computed by Spearman’s rank correlation test with two-tailed p-value and 95% confidence intervals and visualised in ChiPlot.online. All calculations were made in the GraphPad Prism 8.0.1 software (Insight Venture Management, New-York, NY, USA).

## Results

### Expression of *E. coli* PAP-1

To evaluate the production of recombinant PAP-1 in *E. coli* cells, we cloned the PAP-1 coding sequence into the expression vector pET36b, placing it under the control of the lac promoter, thereby generating the plasmid pET36b-PAP1-Eco. The pET36b vector enables tight regulation of recombinant protein expression via the recombinant LacI protein encoded by the plasmid. Additionally, the vector’s antibiotic resistance gene and PAP-1 coding sequence are located in separate transcriptional units, preventing unintended PAP-1 expression due to the synthesis of a polycistronic RNA from the antibiotic resistance gene promoter.

Seven *E. coli* strains were tested for PAP-1 expression, including the commonly used BL21 (DE3) pLysS strain, its derivatives optimized for recombinant protein production, and two K-12 strain derivatives, Rosetta Blue (DE3) and Shuffle T7 (Table 2). This selection allowed us to consider the impact of different genetic backgrounds on PAP-1 expression. After transformation with pET36b-PAP1-Eco, *E. coli* cells were grown to an OD600 of 0.6, and PAP-1 synthesis was induced by the addition of IPTG to a final concentration of 1 mM. The induced cultures were divided into three equal portions and incubated overnight at 18°C, 25°C, and 37°C, respectively. Following induction, the optical density of the overnight cultures was measured, and the relative amount of recombinant PAP-1 was evaluated using SDS-PAGE (Figure 1).

**Figure 1.**
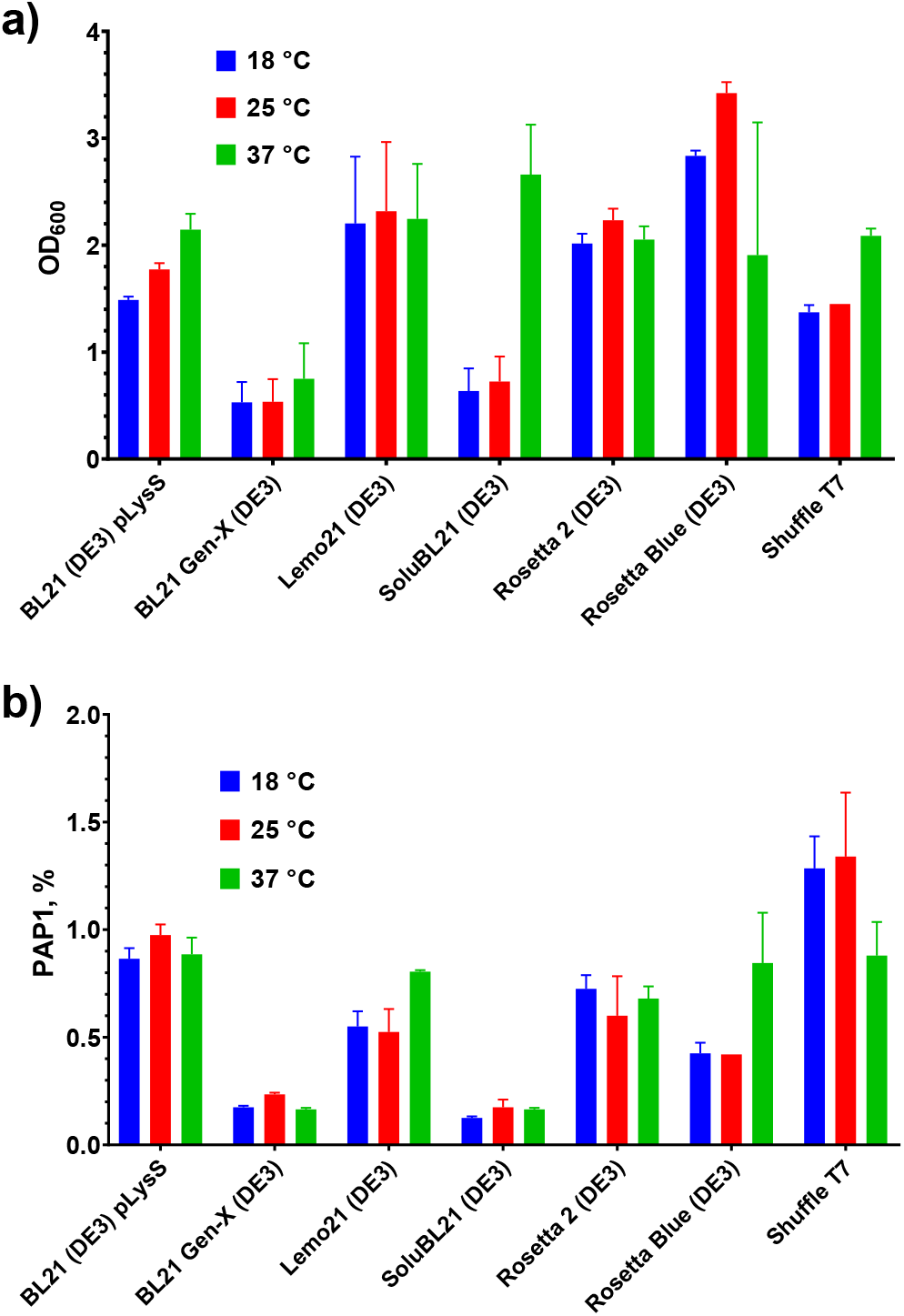
Optical density and level of PAP-1 after PAP-1 expression. a) Optical density of *E. coli* cultures after overnight expression of PAP-1. Y-axis marks OD600, X-axis represents strains. b) Relative quantity of PAP-1. Y-axis designates PAP-1 quantity related to all protein bands, X-axis designates strains. All experiments were triplicated, error bars demonstrate SD.

After PAP-1 induction, the highest optical density (OD600) was observed in the Rosetta Blue (DE3) strain, exceeding 3. In contrast, the BL21 Gen-X strain demonstrated the lowest OD600, which did not exceed 1 across all induction temperatures. Similarly, SoluBL21 showed low growth, except for cultures incubated at 37°C, which achieved an OD600 of 2.6. These findings align with the manufacturer’s notes, which indicate a reduced growth rate for these strains. For the remaining *E. coli* strains, cell density was moderate, with OD600 values ranging from 1.5 to 2.3. Notably, only three strains—BL21 (DE3) pLysS, SoluBL21, and Shuffle T7—exhibited higher cell densities at 37°C compared to other temperatures. For the other strains, OD600 values showed no consistent correlation with induction temperature (Figure 2).

**Figure 2.**
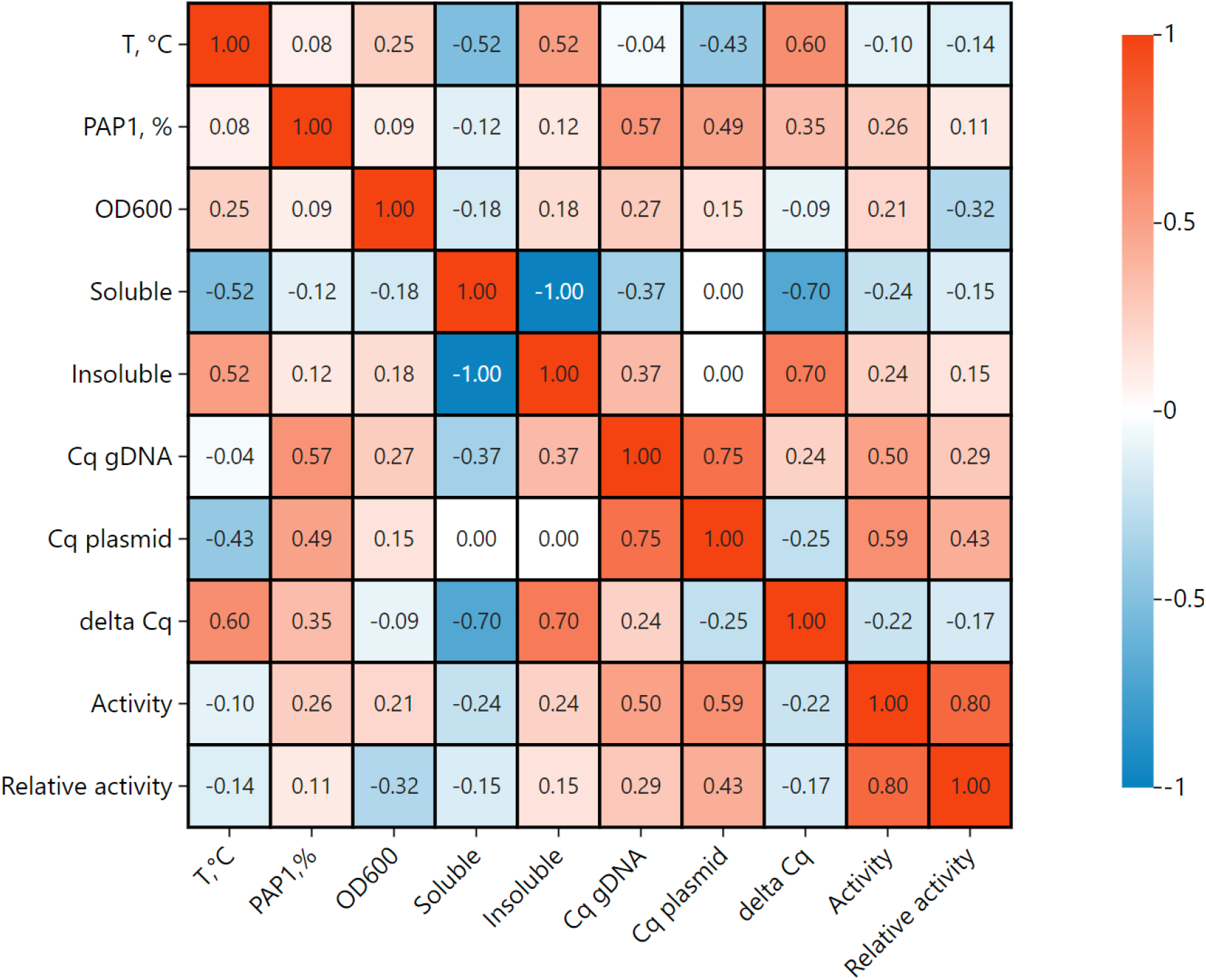
A heatmap of correlation between variables in PAP-1 induction. The heatmap made in ChiPlot (chiplot.online) represents Spearman’s rang coefficients for each pair of variables, the coefficients are indicated in each cell.

The highest PAP-1 levels were observed in the Shuffle T7 and BL21 (DE3) pLysS strains, reaching 1–1.5% of total protein, while the lowest levels were found in BL21 Gen-X and SoluBL21, at 0.18–0.24%. Interestingly, for strains showing increased OD600 at 37°C, PAP-1 levels at this temperature were either lower than or equal to those at 18°C and 25°C. Thus, whether analyzed individually or collectively, PAP-1 levels did not show a positive correlation with cell optical density after induction.

### Solubility of *E. coli* PAP-1

After measuring the bulk PAP-1 level after induction, we assessed solubility of the recombinant protein. For that, the cell pellets were lysed in a buffer containing 200 mM KCl, two detergents (3-(N,N-Dimethylmyristyl-ammonio)propanesulfonate, 0.05% C7BzO) reported as efficient reagents for fast cell disruption and lysozyme. Insoluble and soluble fractions were separated by centrifugation and protein compositions were analyzed using SDS-PAGE (Figure 3).

**Figure 3.**
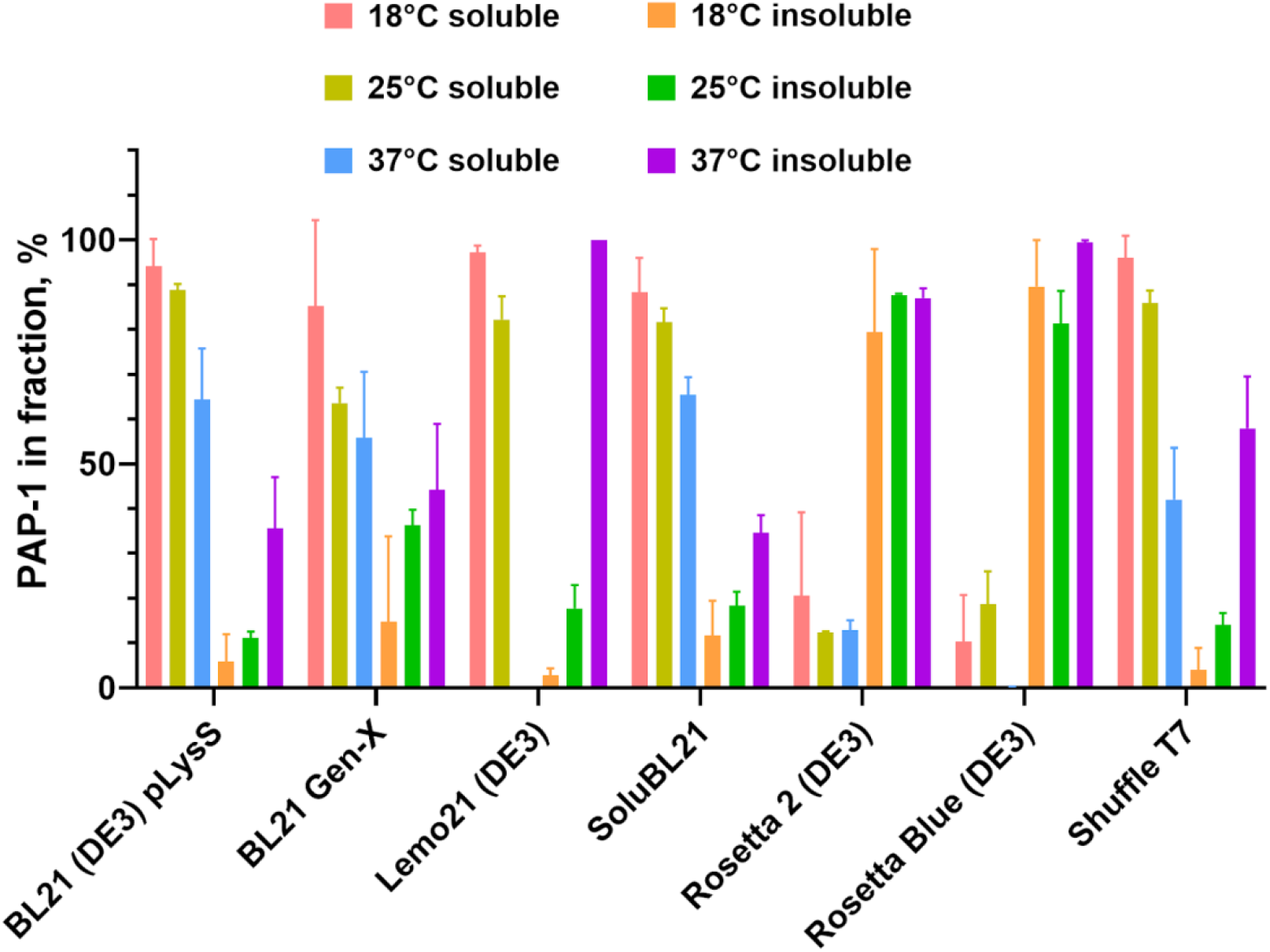
Solubility of PAP-1. Y-axis marks PAP-1 percent in soluble or insoluble fractions, X-axis represents strains. All experiments were triplicated, error bars demonstrate SD.

Unlike the overall PAP-1 levels, the solubility of PAP-1 showed a moderate negative correlation with induction temperature. For all tested *E. coli* strains, higher induction temperatures resulted in decreased solubility. In most strains, PAP-1 was nearly 100% soluble when expressed at 18°C, whereas only 42–65% of the recombinant enzyme remained soluble at 37°C. An exception to this trend was Lemo21 (DE3), where no soluble PAP-1 was detected at 37°C. Additionally, in both Rosetta strains (Rosetta 2 (DE3) and Rosetta Blue (DE3)), PAP-1 solubility was notably lower, with a maximum solubility of approximately 20% at 18°C. These strains originate from different lineages but share the pRARE plasmid, which encodes tRNAs that are rare in *E. coli*.

### Specific activity of *E. coli* PAP-1

Since protein bands on SDS-PAGE do not always accurately reflect the quantity of active enzymes, we further tested poly(A) polymerase activity in the samples after PAP-1 expression. Typically, the endogenous level of PAP-1 in *E. coli* is low, and its associated specific activity is negligible. For this assay, a fluorescently labeled 20-mer A oligonucleotide was incubated with varying volumes of cell lysates in a reaction buffer optimized for *E. coli* PAP-1. To minimize potential inhibition by cellular components, lysate volumes were titrated in the range of 0.015–1 μL per reaction.

Elongated reaction products were quantified, and the lysate volumes required to polyadenylate 50% of the substrate were calculated. These computed lysate volumes served as a measure of PAP-1 activity in each sample. To account for differences in cell density across strains, the calculated lysate volumes were normalized to the respective OD600 values. Higher computed lysate volumes indicated lower PAP-1 activity in the corresponding samples. The results of the activity analysis are presented in Figure 4.

**Figure 4.**
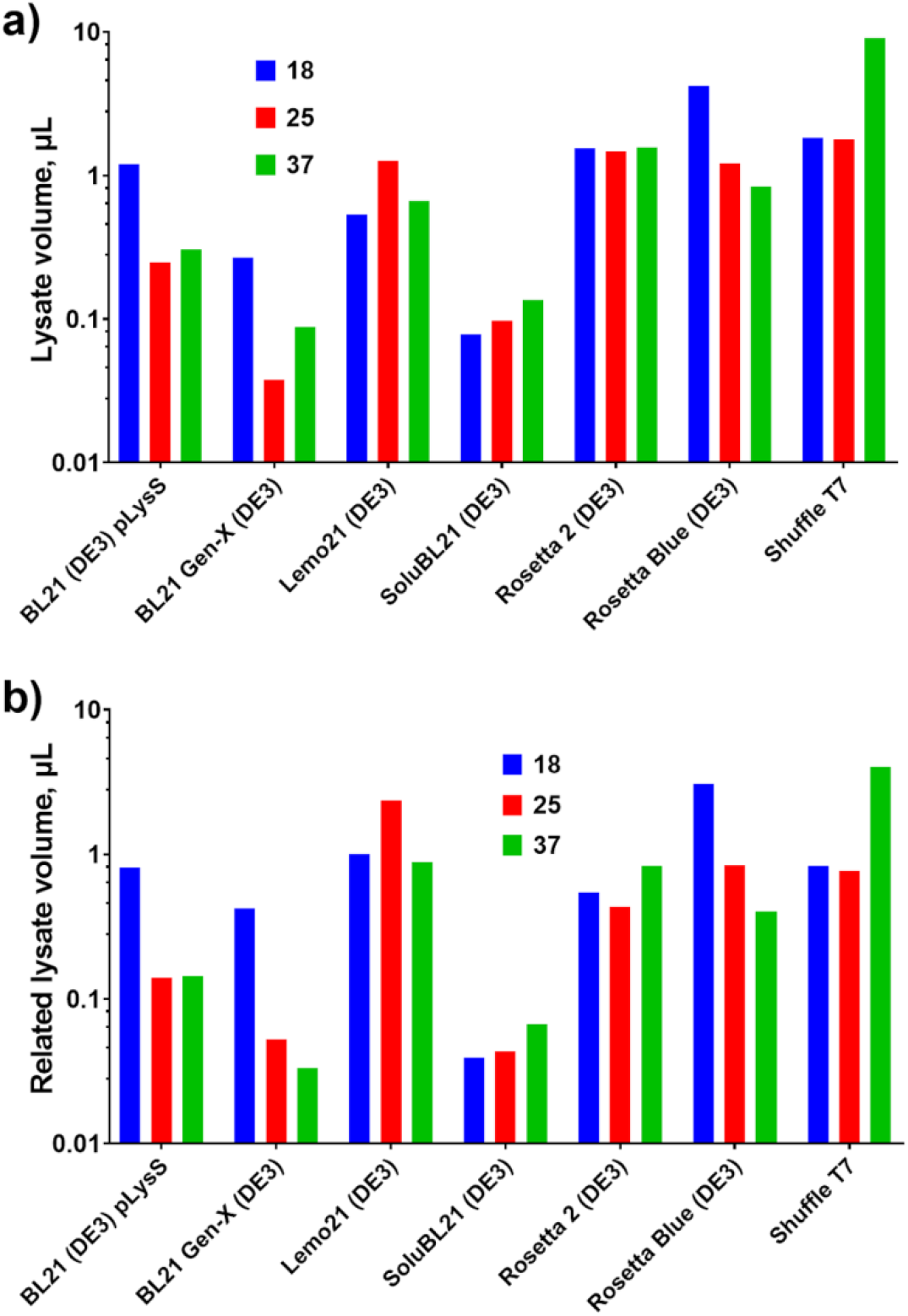
Specific activity of PAP-1. a) Computed lysate volumes for polyadenylation of 50% of the substrate. b) Computed lysate volumes adjusted to the respective OD600. PAP-1 activity was measured using elongation of the fluorescently labeled oligo20(r)A substrate. Y-axis marks percent of the elongated substrate, X-axis represents strains.

A weak positive correlation was observed between PAP-1 activity and PAP-1 solubility: the more PAP-1 remained in the soluble fraction, the more active it was. However, no correlation was found between PAP-1 activity and expression temperature, indicating that PAP-1 remained active even in the insoluble state when the induction temperature was increased. A moderate positive association was noted between PAP-1 activity and the amount of DNA in the cells, whether genomic or plasmid. Notably, increased PAP-1 activity positively correlated with plasmid copy number, suggesting that relaxation of plasmid replication occurred due to the accelerated degradation of antisense RNA-1, which normally inhibits priming at ColE1 origins.

### Relation between plasmid copy number and level of recombinant PAP-1

The PAP-1 gene, pcnB, was identified as a locus involved in the regulation of plasmid numbers in *E. coli* cells. Specifically, PAP-1 polyadenylates a small RNA1 transcript, which negatively regulates the replication of plasmids with a ColE1-based origin. Therefore, we hypothesized that there could be a link between PAP-1 levels and the copy number of the plasmid encoding PAP-1, i.e., high PAP-1 expression might lead to an increase in plasmid numbers in host cells. To test this hypothesis, we measured the amounts of *E. coli* genomic DNA and the pET-PAP-Eco plasmid using qPCR. The results are presented as Cq values in Figure 5a. Notably, Cq values are inversely correlated with the amount of the respective DNA template. Since Cq values are logarithmic, they serve as relative indicators of the DNA template quantity in the analyzed sample.

**Figure 5.**
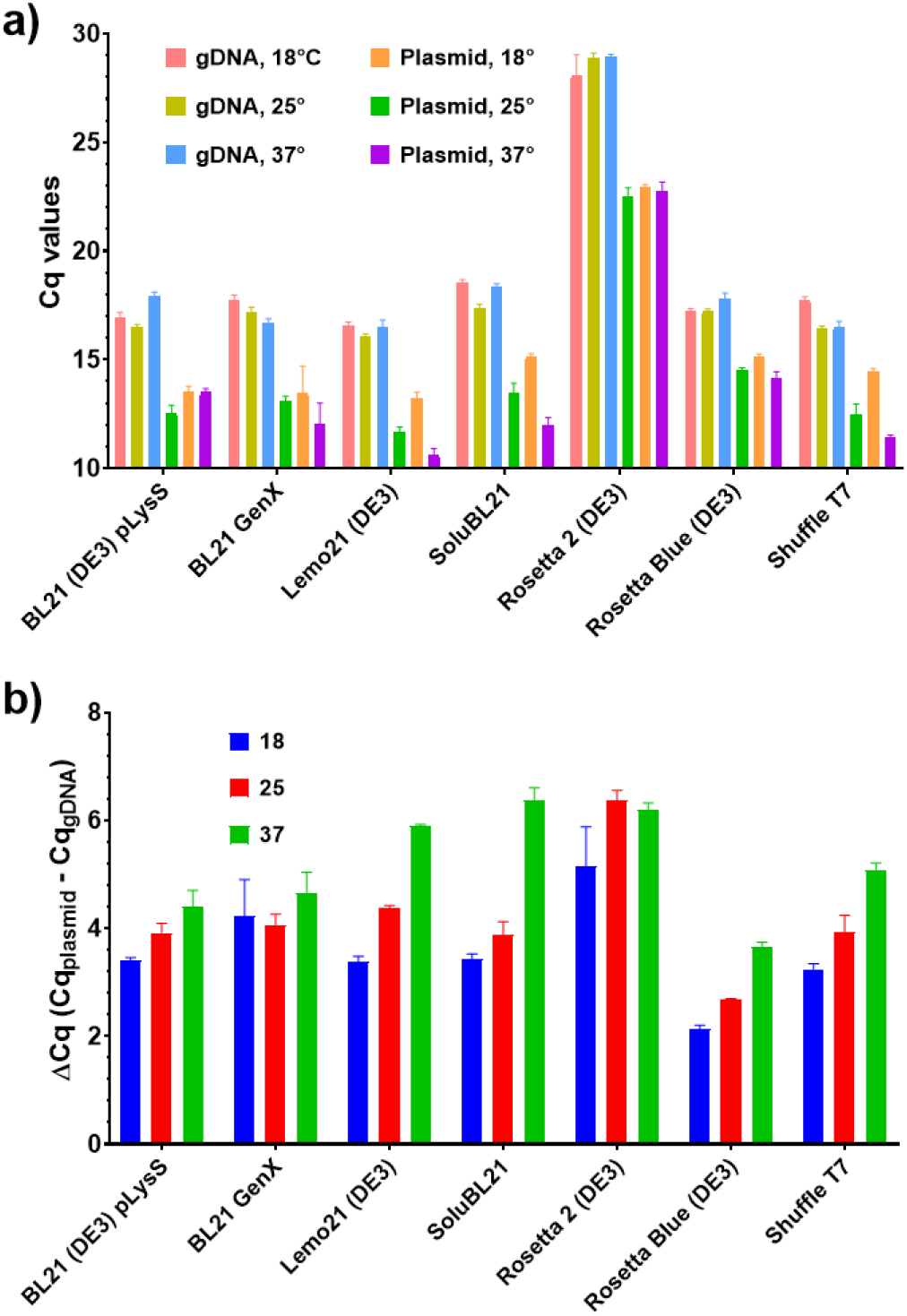
Amount of *E. coli* genomic DNA and PAP-1 encoding plasmid after PAP-1 expression. a) Cq values of qPCR for quantification of *E. coli* genomic DNA and PAP-1 encoding plasmid pET-PAP-Eco. Y-axis marks Cq values, X-axis represents strains. b) Relative quantity of pET- PAP-Eco. Y-axis designates the difference between Cq for the plasmid and gDNA, X-axis designates strains. All experiments were triplicated, error bars demonstrate SD.

Unlike optical density, no correlation was found between expression temperature and Cq values for genomic DNA. Thus, qPCR did not detect any temperature-dependent changes in the accumulation of cellular DNA. However, the situation was different for the pET-PAP-Eco plasmid, where plasmid numbers increased at 37°C compared to 25°C in Lemo21 (DE3), SoluBL21, Rosetta Blue (DE3), and Shuffle T7 strains. In contrast, no significant trend was observed for BL21 Gen-X.

To further investigate potential changes in DNA amounts, we calculated the difference between Cq values for genomic DNA and pET-PAP-Eco plasmid. This difference served as a rough measure of the ratio between the amounts of genomic DNA and the plasmid, assuming similar qPCR efficiency for both targets. The calculated delta Cq values are presented in Figure 5b, showing a moderate positive correlation between expression temperature and pET-PAP-Eco plasmid copy numbers.

Cq values for both genomic DNA and the pET-PAP-Eco plasmid, as well as ΔCq values, were moderately correlated with PAP-1 levels. Surprisingly, a strong negative correlation was observed between ΔCq values and PAP-1 solubility, while solubility was not associated with the plasmid’s Cq values and was positively correlated with the amount of genomic DNA. This suggests that higher insolubility of PAP-1 was associated with higher plasmid copy numbers. However, PAP-1 activity positively correlated with the amounts of genomic DNA, plasmid, and plasmid copy number.

## Discussion

The rapid progress in the development and commercialization of mRNA vaccines has highlighted the importance of enzymes required for mRNA processing, particularly when the enzymatic method is chosen. *E. coli* PAP-1 is one such enzyme, as polyadenylation of mRNA molecules is essential for their stability in eukaryotic cells. Therefore, the efficient production of *E. coli* PAP-1, along with other recombinant poly(A)-polymerases, is a crucial step in mRNA production. However, PAP-1 is toxic to host cells, prone to aggregation, and loses activity when stored in low-salt conditions. In *E. coli*, PAP-1 production is tightly regulated to prevent negative impacts on cell viability and growth rate, making the expression of recombinant PAP-1 a challenging task. Notably, *E. coli* PAP-1 is a difficult protein for recombinant production due to its high pI (9.67), high charge at pH 7.0 (16.51), and the presence of four cysteine residues.

In this study, we examined PAP-1 expression in seven *E. coli* strains known for producing “difficult” recombinant proteins, including toxic, membrane-bound, insoluble proteins, and proteins with a high number of disulfide bonds. BL21 (DE3) pLysS was chosen as a reference strain because it is commonly used for recombinant protein expression. BL21 Gen-X and SoluBL21 (DE3) are optimized for producing mammalian and soluble proteins, respectively. LemoBL21 (DE3) offers tunable expression of recombinant proteins regulated by L-rhamnose concentration. Shuffle T7 is a K-12-derived strain with a DsbC chaperone to correct non-native disulfide bonds and additional LacI to prevent lac promoter leakage. Rosetta 2 (DE3) is a BL21 (DE3) derivative with a pRARE2 plasmid, providing tRNAs to suppress codon bias for eukaryotic proteins. Rosetta Blue (DE3) is a K-12-based strain with the same pRARE plasmid and a lacIq mutation, similar to Shuffle T7. Thus, five strains were from the B lineage of *E. coli*, while two originated from the K- 12 lineage. The B-lineage strains lack the cytoplasmic Lon and membrane OmpT proteases, but it was unknown whether these enzymatic activities affect PAP-1 production in *E. coli*. However, no differences in optical density, relative amount, or solubility were observed between the B- and K-12 strains.

Previous studies have shown that *E. coli* PAP-1 is highly toxic, causing cell death within 30 minutes of IPTG induction [6]. However, in our experiment, all strains except for the slow-growing BL21 Gen-X (DE3) and SoluBL21 (DE3) reached an OD600 of 1.5–3.5, and PAP-1 production was directly associated with final optical density. While we did not assess cell viability post-induction, these results seem to contradict the reported toxicity of PAP-1. This discrepancy may be explained by the delayed toxic effects of PAP-1, where high enzyme production does not cause immediate cell death.

Both Rosetta strains, derived from different lineages but sharing the pRARE plasmid, exhibited the lowest solubility of PAP-1. It remains unclear whether the pRARE plasmid or other unknown factors contributed to PAP-1 aggregation into inclusion bodies. One possible explanation is that the higher translation rate of PAP-1, facilitated by the rare tRNAs encoded by the pRARE plasmid, may lead to aggregation. This hypothesis is supported by several observations: first, rare codons generally have a minimal impact on protein expression in *E. coli* [32]; second, an increased translation rate of heterologous proteins in *E. coli* strains with the pRIL plasmid encoding rare tRNAs has been associated with reduced solubility [33]; and third, a high percentage of arginine correlates with lower expression and solubility [34]. The nucleotide sequence of PAP-1 contains 18 rare codons (mainly encoding threonine) per 466 codons, as well as an unusually high proportion of arginine residues (11.4%, or 53 residues). However, we did not directly test this hypothesis by comparing PAP-1 expression with and without the pRARE plasmid. In other strains, solubility depended on the expression temperature without any notable anomalies.

The activity and level of PAP-1 were lowest in BL21 Gen-X (DE3) and SoluBL21 (DE3), the slowest-growing strains. We speculated that the low PAP-1 production in these strains was due to its toxicity. Other strains, which grew faster, produced more cells expressing recombinant PAP-1. Therefore, if these *E. coli* cells lost viability due to PAP-1 accumulation, many of the dead cells would still contain the active enzyme. Conversely, slower-growing strains did not produce a substantial quantity of PAP-1 due to limited time for expression, resulting in a less productive culture.

The lack of association between solubility and the specific activity of PAP-1 can be attributed to several potential explanations. First, the enzyme may retain its activity even in an insoluble state. Second, PAP-1 might gradually dissolve from inclusion bodies when exposed to the high ionic strength of the PAP-1 reaction buffer. This second hypothesis is supported by PAP- 1’s known tendency to aggregate under low-salt conditions. However, whether these assumptions are valid or if other factors influence PAP-1 activity remains unclear.

Intriguing results emerged when we explored the potential correlations between DNA amounts, plasmid copy numbers, and PAP-1 properties. Higher PAP-1 activity corresponded with an increased plasmid DNA quantity and a higher plasmid copy number. This observation aligns with PAP-1’s role in degrading RNA-1, which negatively regulates replication of ColE1-based plasmids [4]. However, the strong negative association between solubility and plasmid copy numbers was unexpected. We speculate that the rapid accumulation of PAP-1 facilitated efficient plasmid propagation, leading to its aggregation and incorporation into inclusion bodies. Thus, despite the relatively low solubility of PAP-1 in Rosetta strains, its activity was still evident through changes in plasmid copy numbers.

The increased genomic DNA content observed presented another challenge, as no direct link between PAP-1, polyadenylation, and *E. coli* genome replication has been identified. This elevated DNA content may result from other cellular processes not accounted for in this study.

To summarize, we evaluated the ability of several *E. coli* strains to produce recombinant *E. coli* PAP-1 at different induction temperatures by analyzing the enzyme’s overall level, solubility, and specific activity.

## Conclusion

In this study, we evaluated the production of recombinant PAP-1 across seven different *E. coli* strains. The BL21 (DE3) pLysS strain demonstrated the optimal balance of cell density, overall PAP-1 yield, solubility, and specific activity. In contrast, Rosetta strains displayed the lowest solubility of the recombinant enzyme, potentially due to excessively high translation efficiency. Induction temperatures above 18°C led to increased insolubility of PAP-1. Notably, PAP-1 accumulation was associated with an increase in the plasmid copy number encoding the enzyme, suggesting its potential use as a surrogate marker for PAP-1 activity.

## CRediT

**Igor P. Oscorbin**: Methodology, Validation, Data Curation, Writing - Original Draft, Supervision. **Maria S. Kunova**: Investigation, Visualization. **Maxim L. Filipenko**: Conceptualization, Writing - Review & Editing.

## Acknowledgements

This research and the APC were funded by Russian Science Foundation, grant number 24-24- 00389, https://www.rscf.ru/project/24-24-00389.

## References

1. Hajnsdorf, E.; Kaberdin, V.R. RNA Polyadenylation and Its Consequences in Prokaryotes. Philos. Trans. R. Soc. B Biol. Sci. 2018, 373, 20180166, doi:10.1098/rstb.2018.0166.

2. Mohanty, B.K.; Kushner, S.R. Polynucleotide Phosphorylase Functions Both as a 3′ ? 5′ Exonuclease and a Poly(A) Polymerase in Escherichia Coli. Proc. Natl. Acad. Sci. 2000, 97, 11966– 11971, doi:10.1073/pnas.220295997.

3. Bralley, P.; Cozad, M.; Jones, G.H. Geobacter Sulfurreducens Contains Separate C- and A-Adding TRNA Nucleotidyltransferases and a Poly(A) Polymerase. J. Bacteriol. 2009, 191, 109–114, doi:10.1128/JB.01166-08.

4. Lopilato, J.; Bortner, S.; Beckwith, J. Mutations in a New Chromosomal Gene of Escherichia Coli K-12, PcnB, Reduce Plasmid Copy Number of PBR322 and Its Derivatives. Mol. Gen. Genet. MGG 1986, 205, 285–290, doi:10.1007/BF00430440.

5. Cao, G.J.; Sarkar, N. Identification of the Gene for an Escherichia Coli Poly(A) Polymerase. Proc. Natl. Acad. Sci. 1992, 89, 10380–10384, doi:10.1073/pnas.89.21.10380.

6. Mohanty, B.K.; Kushner, S.R. Analysis of the Function of Escherichia Coli Poly(A) Polymerase I in RNA Metabolism. Mol. Microbiol. 1999, 34, 1094–1108, doi:10.1046/j.1365-2958.1999.01673.x.

7. Maes, A.; Gracia, C.; Innocenti, N.; Zhang, K.; Aurell, E.; Hajnsdorf, E. Landscape of RNA Polyadenylation in E. Coli. Nucleic Acids Res. 2016, gkw894, doi:10.1093/nar/gkw894.

8. Jones, G.H. Phylogeny and Evolution of RNA 3′-Nucleotidyltransferases in Bacteria. J. Mol. Evol. 2019, 87, 254–270, doi:10.1007/s00239-019-09907-2.

9. August, J.T.; Ortiz, P.J.; Hurwitz, J. Ribonucleic Acid-Dependent Ribonucleotide Incorporation. I. Purification and Properties of the Enzyme. J. Biol. Chem. 1962, 237, 3786–3793.

10. Kalapos, M.P.; Cao, G.J.; Kushner, S.R.; Sarkar, N. Identification of a Second Poly(A) Polymerase in Escherichia Coli. Biochem. Biophys. Res. Commun. 1994, 198, 459–465, doi:10.1006/bbrc.1994.1067.

11. Bralley, P.; Cozad, M.; Jones, G.H. Geobacter Sulfurreducens Contains Separate C- and A-Adding TRNA Nucleotidyltransferases and a Poly(A) Polymerase. J. Bacteriol. 2009, 191, 109–114, doi:10.1128/JB.01166-08.

12. Payne, K.J.; Boezi, J.A. The Purification and Characterization of Adenosine Triphosphate Ribonucleic Acid Adenyltransferase from Pseudomonas Putida. J. Biol. Chem. 1970, 245, 1378– 1387.

13. Sippel, A.E. Purification and Characterization of Adenosine Triphosphate: Ribonucleic Acid Adenyltransferase from Escherichia Coli. Eur. J. Biochem. 1973, 37, 31–40, doi:10.1111/j.1432-1033.1973.tb02953.x.

14. Yehudai-Resheff, S. Characterization of the E.Coli Poly(A) Polymerase: Nucleotide Specificity, RNA-Binding Affinities and RNA Structure Dependence. Nucleic Acids Res. 2000, 28, 1139–1144, doi:10.1093/nar/28.5.1139.

15. Liu, J.D.; Parkinson, J.S. Genetics and Sequence Analysis of the PcnB Locus, an Escherichia Coli Gene Involved in Plasmid Copy Number Control. J. Bacteriol. 1989, 171, 1254–1261, doi:10.1128/jb.171.3.1254-1261.1989.

16. Jones, G.H. Acquisition of PcnB [Poly(A) Polymerase I] Genes via Horizontal Transfer from the β, γ-Proteobacteria. Microb. Genomics 2021, 7, doi:10.1099/mgen.0.000508.

17. Chaudhary, N.; Weissman, D.; Whitehead, K.A. MRNA Vaccines for Infectious Diseases: Principles, Delivery and Clinical Translation. Nat. Rev. Drug Discov. 2021, 20, 817–838, doi:10.1038/s41573-021-00283-5.

18. Sayour, E.J.; Boczkowski, D.; Mitchell, D.A.; Nair, S.K. Cancer MRNA Vaccines: Clinical Advances and Future Opportunities. Nat. Rev. Clin. Oncol. 2024, 21, 489–500, doi:10.1038/s41571-024-00902-1.

19. Rosano, G.L.; Morales, E.S.; Ceccarelli, E.A. New Tools for Recombinant Protein Production in Escherichia Coli : A 5-year Update. Protein Sci. 2019, 28, 1412–1422, doi:10.1002/pro.3668.

20. İncir, İ.; Kaplan, Ö. Escherichia Coli as a Versatile Cell Factory: Advances and Challenges in Recombinant Protein Production. Protein Expr. Purif. 2024, 219, 106463, doi:10.1016/j.pep.2024.106463.

21. Tegel, H.; Tourle, S.; Ottosson, J.; Persson, A. Increased Levels of Recombinant Human Proteins with the Escherichia Coli Strain Rosetta(DE3). Protein Expr. Purif. 2010, 69, 159–167, doi:10.1016/j.pep.2009.08.017.

22. Redda, Y.T.; Venkatesh, G.; Kalaiyarasu, S.; Bhatia, S.; Kumar, D.S.; Nagarajan, S.; Pillai, A.; Tripathi, S.; Kulkarni, D.D.; Dubey, S.C. Expression and Purification of Recombinant H5HA1 Protein of H5N1 Avian Influenza Virus in E. Coli and Its Application in Indirect ELISA. J. Immunoass. Immunochem. 2016, 37, 346–358, doi:10.1080/15321819.2015.1135160.

23. Ossysek, K.; Uchański, T.; Kulesza, M.; Bzowska, M.; Klaus, T.; Woś, K.; Madej, M.; Bereta, J. A New Expression Vector Facilitating Production and Functional Analysis of ScFv Antibody Fragments Selected from Tomlinson I+J Phagemid Libraries. Immunol. Lett. 2015, 167, 95–102, doi:10.1016/j.imlet.2015.07.005.

24. Huynh Thi Yen, L.; Park, S.-Y.; Kim, J.-S. Cloning, Crystallization and Preliminary X-Ray Diffraction Analysis of an Intact DNA Methyltransferase of a Type I Restriction-Modification Enzyme from Vibrio Vulnificus. Acta Crystallogr. Sect. F, Struct. Biol. Commun. 2014, 70, 489–492, doi:10.1107/S2053230X14004543.

25. Hata, S.; Kitamura, F.; Sorimachi, H. Efficient Expression and Purification of Recombinant Human μ-Calpain Using an Escherichia Coli Expression System. Genes Cells 2013, 18, 753–763, doi:10.1111/gtc.12071.

26. Yamaguchi, R.; Akter, S.; Kanehama, A.; Iwamoto, T.; Hasegawa, M.; Ito, A.; Nishimukai, M.; Yamada, M.; Kashiwagi, A. Improvement of Solubility of Phospholipase D from Streptomyces Antibioticus in Recombinant Escherichia Coli and Its Application for the Enzymatic Synthesis of a Non-Natural Plasmalogen. Lett. Appl. Microbiol. 2023, 76, doi:10.1093/lambio/ovad049.

27. Diankristanti, P.A.; Effendi, S.S.W.; Hsiang, C.-C.; Ng, I.-S. High-Level Itaconic Acid (IA) Production Using Engineered Escherichia Coli Lemo21(DE3) toward Sustainable Biorefinery. Enzyme Microb. Technol. 2023, 167, 110231, doi:10.1016/j.enzmictec.2023.110231.

28. Schlegel, S.; Löfblom, J.; Lee, C.; Hjelm, A.; Klepsch, M.; Strous, M.; Drew, D.; Slotboom, D.J.; de Gier, J.-W. Optimizing Membrane Protein Overexpression in the Escherichia Coli Strain Lemo21(DE3). J. Mol. Biol. 2012, 423, 648–659, doi:10.1016/j.jmb.2012.07.019.

29. Nasiri, M.; Babaie, J.; Amiri, S.; Azimi, E.; Shamshiri, S.; Khalaj, V.; Golkar, M.; Fard-Esfahani, P. SHuffleTM T7 Strain Is Capable of Producing High Amount of Recombinant Human Fibroblast Growth Factor-1 (RhFGF-1) with Proper Physicochemical and Biological Properties. J. Biotechnol. 2017, 259, 30–38, doi:10.1016/j.jbiotec.2017.08.015.

30. Lobstein, J.; Emrich, C.A.; Jeans, C.; Faulkner, M.; Riggs, P.; Berkmen, M. SHuffle, a Novel Escherichia Coli Protein Expression Strain Capable of Correctly Folding Disulfide Bonded Proteins in Its Cytoplasm. Microb. Cell Fact. 2012, 11, 56, doi:10.1186/1475-2859-11-56.

31. Evans, G.A. Molecular Cloning: A Laboratory Manual. Second Edition. Volumes 1, 2, and 3. Current Protocols in Molecular Biology. Volumes 1 and 2; Cold Spring Harbor Laboratory Press: New York, 1990; Vol. 61; ISBN 13: 978-0879693091.

32. Boël, G.; Letso, R.; Neely, H.; Price, W.N.; Wong, K.-H.; Su, M.; Luff, J.D.; Valecha, M.; Everett, J.K.; Acton, T.B.; et al. Codon Influence on Protein Expression in E. Coli Correlates with MRNA Levels. Nature 2016, 529, 358–363, doi:10.1038/nature16509.

33. Rosano, G.L.; Ceccarelli, E.A. Rare Codon Content Affects the Solubility of Recombinant Proteins in a Codon Bias-Adjusted Escherichia Coli Strain. Microb. Cell Fact. 2009, 8, 41, doi:10.1186/1475-2859-8-41.

34. Price, W.N.; Handelman, S.K.; Everett, J.K.; Tong, S.N.; Bracic, A.; Luff, J.D.; Naumov, V.; Acton, T.; Manor, P.; Xiao, R.; et al. Large-Scale Experimental Studies Show Unexpected Amino Acid Effects on Protein Expression and Solubility in Vivo in E. Coli. Microb. Inform. Exp. 2011, 1, 6, doi:10.1186/2042-5783-1-6.

